# The Healthy Brain Network Serial Scanning Initiative: A resource for evaluating inter-individual differences and their reliabilities across scan conditions and sessions

**DOI:** 10.1101/078881

**Authors:** David O’Connor, Natan Vega Potler, Meagan Kovacs, Ting Xu, Lei Ai, John Pellman, Tamara Vanderwal, Lucas Parra, Samantha Cohen, Satrajit Ghosh, Jasmine Escalera, Natalie Grant-Villegas, Yael Osman, Anastasia Bui, R. Cameron Craddock, Michael P. Milham

**Author notes:** Correspondence: Michael Peter Milham, MD, PhD, Center for Developing Brain, Child Mind Institute, New York, NY 10022, USA (M.P. Milham).

## Abstract

**Background:** Although typically measured during the resting state, a growing literature is illustrating the ability to map intrinsic connectivity in task and naturalistic viewing fMRI paradigms. These paradigms are drawing excitement due to their greater tolerability in clinical and developing populations and because they enable a wider range of analyses (e.g. inter-subject correlations). To be clinically useful, the test-retest reliability of connectivity measured during these paradigms needs to be established. This resource provides data for evaluating testretest reliability for full-brain connectivity patterns detected during each of four scan conditions that differ with respect to level of engagement (rest, abstract animations, movie clips, flanker task). Data is provided for thirteen participants, each scanned in twelve sessions with 10 minutes for each scan of the four conditions. Diffusion kurtosis imaging data was also obtained at each session.

**Findings:** Technical validation and demonstrative reliability analyses found that variation in intrinsic functional connectivity across sessions was greater than that attributable to scan condition. Between-condition reliability was generally high, particularly for the frontoparietal and default networks. Between-session reliabilities obtained separately for the different scan conditions were comparable, though notably lower than between-condition reliabilities.

**Conclusions:** The described resource provides a test-bed for quantifying the reliability of connectivity indices across conditions and time. The resource can be used to compare and optimize different frameworks for measuring connectivity and data collection parameters such as scan length. Additionally, investigators can explore the unique perspectives of the brain’s functional architecture offered by each of the scan conditions.

## DATA NOTE

### Purpose of Data Collection

An extensive literature has documented the utility of fMRI for mapping the brain’s functional interactions through the detection of temporally correlated patterns of spontaneous activity between spatially distinct brain areas [1]–[7]. Commonly referred to as intrinsic functional connectivity (iFC), these patterns are commonly studied during the ‘resting state’, which involves the participant quietly lying awake and not performing an externally driven task. Resting state fMRI (R-fMRI) has gained popularity in clinical neuroimaging due to its minimal task and participant compliance demands. R-fMRI has also demonstrated good test-retest reliability for commonly used measures [8]–[12], and utility in detecting brain differences associated with neuropsychiatric disorders [13], [14]. Despite these successes, a growing body of work is questioning the advantages of resting state, given reports of higher head motion, decreased tolerance of the scan environment (e.g. boredom, rumination), and increased likelihood of falling asleep compared to more engaging task-based fMRI paradigms [15]–[18]. This is particularly relevant for studies of pediatric, geriatric and clinical populations, all of which are characterized by lower tolerance of the scanner environment.

A number of less challenging scan conditions have been proposed as alternatives for estimating iFC. Particularly intriguing are “naturalistic viewing” paradigms [15], [19], [20]. It has been shown that the mental state (i.e., emotional state, performing a task, etc.) of the participant during scanning can effect iFC patterns; recent work suggests that low engagement states (e.g., computer animations with limited cognitive content) may come close to mimicking rest from a neural perspective [21]. Several studies have illustrated the ability to relate trait phenotypic variables to inter-individual differences in iFC across conditions, even if extrinsically driven signals (i.e., task stimulus functions) are not removed [21]–[27]. However, comprehensive comparisons of the relative impact of scan condition on detection of inter-individual differences in intrinsic functional connectivity, and the test-retest reliability of these differences, are needed before these paradigms can fully supplant R-fMRI.

Here we describe a dataset that was generated as part of a pilot testing effort for the Child Mind Institute Healthy Brain Network – a large-scale data collection effort focused on the generation of an open resource for studying child and adolescent mental health. The primary goal of the data collection was to assess and compare test-retest reliability of fullbrain connectivity patterns detected for each of four scan conditions that differed with respect to level of engagement. Specifically, 13 participants were scanned during each of the following four conditions on 12 different occasions: 1) rest, 2) free viewing of abstract computer graphics and sounds designed to have minimal cognitive or emotional content (i.e., “Inscapes”, [15]), 3) free viewing of highly engaging movies [19], and 4) performance of an active task (i.e., an Erickson flanker task [28], with no-Go trials included). For each of the non-rest conditions, three different stimuli were used, with each being repeated four times across the 12 sessions to enable the evaluation of repetition effects. Given the focus on naturalistic viewing, an additional scan session containing a full viewing of “Raiders of the Lost Ark” was included to facilitate interested parties in the exploration and evaluation of increasingly popular hyper alignment approaches, which offer unique solutions to matching brain function across individuals [29].

Although not a primary focus of the data collection, additional structural imaging data was collected, which are being shared as well: 1) MPRAGE [30], 2) diffusion kurtosis imaging [31], [32], 3) quantitative T1/T2 anatomical imaging (single session) [33], 4) magnetization transfer (single session) [34] (see Table 1). Functional MRI data from a single movie viewing session during which Raiders of the Lost Arc was viewed in its entirety, is included as well.

**Table 1.** HBN-SSI experimental design.

| Shared Imaging Data |  |  |
| --- | --- | --- |
| Session # | Session Type | Description |
| 1 | Baseline Characterization | •Multiecho MPRAGE<br>•Diffusion Kurtosis Imaging<br>•Quantitative T1/T2 Mapping<br>•Myelin Transfer Ratio<br>•FLAIR<br>•fMRI: rest (10 min) |
| 2-7, 9-14 | Repeat Scanning | •Multiecho MPRAGE<br>•Diffusion Kurtosis Imaging<br>•fMRI: rest (10 min)<br>•fMRI: Naturalistic Viewing: Inscapes (10 min)<br>•fMRI: Naturalistic Viewing: Movie Clips (10 min)<br>•fMRI: Flanker Task (10 min) |
| 8 | Full Feature Movie | •fMRI: Raiders of the Lost Arc (20 min X 6) |

## METHODS

### Participants and Procedures

13 adults (ages 18-45 years; mean age: 30.3; 38.4% male) recruited from the community participated in the Healthy Brain Network’s Serial Scanning Initiative. Each participant attended 14 sessions over a period of 1-2 months; see Table 1 for the breakdown of data acquired across sessions. All imaging data were collected using a 1.5T Siemens Avanto equipped with a 32-channel head coil in a mobile trailer (Medical Coaches, Oneonta, NY). The scanner was selected as part of a pilot initiative being carried out to evaluate the capabilities of a 1.5T mobile scanner when equipped with a state-of-the-art head coil and imaging sequences. All research performed was approved by the Chesapeake Institutional Review Board, Columbia, MD (https://www.chesapeakeirb.com/).

### Experimental Design

As outlined in Table 1, each participant attended a total of 14 separate imaging session; these included: 1) a baseline characterization session containing a variety of quantitative anatomical scans, 2) 12 serial scanning sessions, each using the same imaging protocol consisting of four functional MRI scan conditions (10 minutes per condition), diffusion kurtosis imaging and a reference MPRAGE anatomical scan, and 3) a ‘Raiders of the Lost Arc’ movie viewing session.

#### Functional MRI Scan Conditions Included in Serial Scanning

The following four functional scan conditions were selected to sample a range of levels of engagement, presented **in ascending order of level of engagement** (**See** Figure 1):

**Figure 1.**
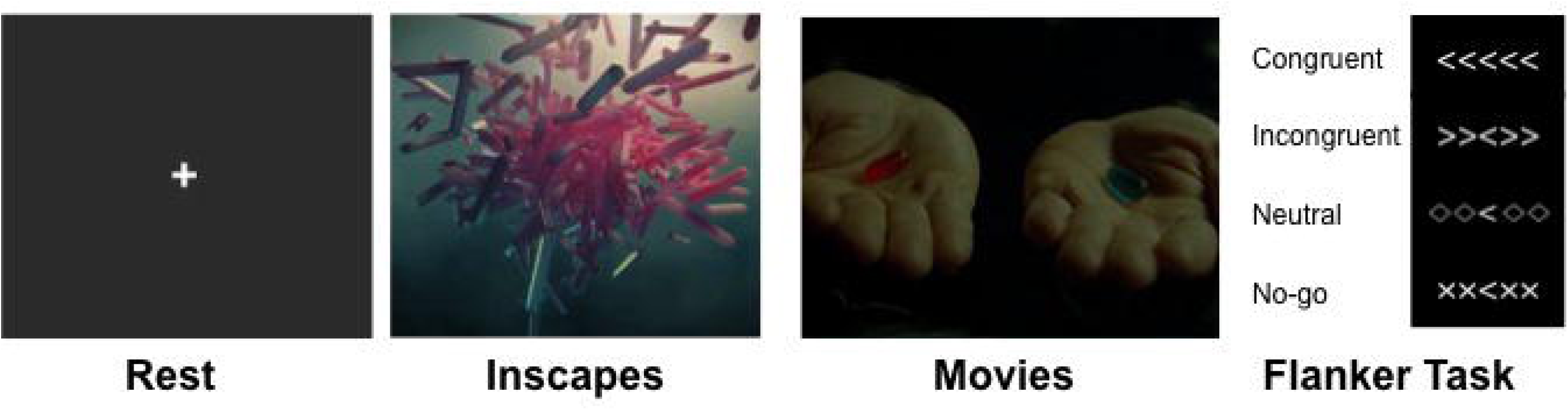
Shown here are sample stimuli from each of the four scan conditions included in the present work. These included: 1) Resting State, (far left), 2) Inscapes (middle left), 3) Movie Clips (e.g., the Matrix; middle right), and 4) Flanker Task (with no-go trials).

#### Rest

The participant was presented a white fixation cross in the center of a black screen and instructed to rest with eyes open. Specific instructions were as follows: “Please lie quietly with your eyes open, and direct your gaze towards the plus symbol. During this scan let your mind wander. If you notice yourself focusing on a particular stream of thoughts, let your mind wander away.”

#### Inscapes

Inscape is a computer generated animation comprised of abstract, non-social, technological-looking 3D forms that transition in a continuous fashion without scene cuts. Visual stimulation is accompanied by repetitive, slow tempo (48 bpm) music based on the pentatonic scale, which was previously selected based on calming influences and to harmonize with the noise generated by EPI sequences [15]. Three unique 10 minute Inscapes were presented across the 12 repeat scanning sessions.

#### Movie

Three unique 10-minute movie clips were presented across the 12 repeat scanning sessions. To ensure a high level of engagement, three Hollywood movie clips were selected, each representing a different movie genre. The specific clips selected were: Wall-E (time codes 00:02:03:13 to 00:12:11:05), The Matrix (00:25:23:10 to 00:35:19:20), and A Few Good Men (01:58:13:01 to 02:08:11:18).

#### Flanker

The Eriksen Flanker task consisted of presenting a series of images containing 5 arrows. For each image, the participant was asked to focus on the center arrow and indicate if it is pointing left or right by pushing a button with their left or right index finger. The flanking arrows could be pointing the same way (congruent) or the opposite way (incongruent). Also built into the task were a neutral stimulus and a go/no-go aspect. The neutral task would contain diamonds instead of flanking arrows, making the central arrow direction more obvious. The no-go stimuli contains x’s instead of flanking arrows, indicating the subject should not push either button. See Figure 1 for a visualization of the stimuli.

#### Counter-Balancing

Order effects are an obvious concern when comparing the four functional scan conditions. To minimize these effects, we ensured that for each participant; 1) each scan type occurred an equal number of times in each of the four scan slots across the 12 sessions, and that 2) each scan type had an equal frequency of being preceded by each of the other three scan types. We made use of 3 exemplars of each non-rest stimuli to enable the examination of repetition effects. For movies, this involved having three 10-minute clips, each from a different movie; for inscapes, this involved three different animation sequences and for the flanker task, three different stimulus orderings were used. We guaranteed that across the 12 scan sessions, each exemplar occurred one time across every three scan sessions. Specific ordering of exemplars were varied across ‘odd’ and ‘even’ numbered participants. For each participant, individual-specific ordering information is provided in the release.

### Imaging Protocols (See Table 2 for scan protocol details)

**Table 2.** MRI acquisition parameters for scans included in the HBN-SSI.

| Image | Whole Brain T1 | Inversion Recovery | Whole Brain T2 | Magnetization Transfer | ME-MPRAGE RMS | T2 FLAIR | DWI | DKI | Rest/Movie/Inscapes/Flanker |
| --- | --- | --- | --- | --- | --- | --- | --- | --- | --- |
| Manufacturer | Siemens |  |  |  |  |  |  |  |  |
| Model | Avanto |  |  |  |  |  |  |  |  |
| Head Coil | 32 Channel |  |  |  |  |  |  |  |  |
| Field Strength | 1.5T |  |  |  |  |  |  |  |  |
| Sequence | 3D Despot 1 | IR-SPGR / 3D Despot 1 | 3D Despot 2 | 3D FLASH | ME-MPRAGE/ 3D TFL | FLAIR | EPI | EPI | EPI |
| Flip Angle | 16 | 5 | 60 | 15 | 7 | 150 | 90 | 90 | 55 |
| Phase Cycling | ? | ? | ? | ? | NA | NA | NA | NA | NA |
| Inversion Time | NA | 400 | NA | NA | 1000 | 2500 | NA | NA | NA |
| Echo Time | 2.4 | 2.4 | 2.7 | 11 | 1.64 | 89 | 76.2 | 93.8 | 40 |
| Repitition Time | 5.2 | 5.3 | 5.4 | 30 | 2730 | 9000 | 3110 | 4500 | 1450 |
| Bandwidth per Voxel (Readout) | 350 | 350 | 350 | 350 | 651 | 190 | 1628 | 1628 | 2374 |
| Parallel Acquisition | None | None | None | GP2 | GP2 | GP2 | MB3 | None | MB3 |
| Partial Fourier | P6/8 S7/8 | S6/8 | P6/8 S7/8 | P6/8 S6/8 | None | None | P6/8 | P6/8 | None |
| Slice Orientation | S | S | S | S | S | T | T | T | T |
| Slice Phase Encoding Direction | AP | AP | AP | AP | AP | RL | AP/PA | AP | AP |
| Slice Acquisition Order | SA | SA | SA | IA | IA | IA | IA | IA | IA |
| Slice Gap | 0.2 | 0.2 | 0.2 | 0.2 | 50 | 0.3 | 0 | 0 | 2.5 |
| Field of View | 220x220 | 220x220 | 220x220 | 256x256 | 256x256 | 201x230 | 192x192 | 192x192 | 192x192 |
| Reconstructed Image Matrix | 128x128x96 | 128x128x48 | 128x128x96 | 256x256x176 | 256x256x176 | 448x512x25 | 96x96x72 | 96x96x72 | 78x78x54 |
| Reconstructed Resolution | 1.72x1.72x1.8 | 1.72x1.72x3.6 | 1.72x1.72x1.8 | 1.0x1.0 | 1.0x1.0x1.0 | 0.45x0.45x6.5 | 2.0x2.0x2.0 | 2.0x2.0x2.0 | 2.46x2.46x2.5 |
| Number of Measurements | 8 | 1 | 16 | 1 | 4 | 1 | 1 | 1 | 420 |
| Acqisition Time | 5;00 | 0;53 | 8;38 | 6;41 | 6;32 | 2;44 | 0;16 | 9;59 | 10;18 |
| Fat Supression | None | None | None | None | None | On | On | On | On |
| Number of Directions | NA | NA | NA | NA | NA | NA | 64 | 64 | NA |
| Number of B Zeros | NA | NA | NA | NA | NA | NA | 1 | 1 | NA |
| B Value (s) | NA | NA | NA | NA | NA | NA | 0 | 0;1000;2000 | NA |
| Averages | NA | NA | NA | NA | 1 | NA | NA | ? | NA |

Legend: AP: Anterior Posterior, PA: Posterior Anterior, RL: Right Left, IA: Interleaved Ascending, SA: Sequential Ascending, S: Saggital, T: Transverse

- Functional MRI (sessions 1-14) For all functional MRI scans, the multiband EPI sequence provided by CMRR [35] was employed to provide high spatial and temporal resolutions (multiband factor 3, voxel size: 2.46x2.46x2.5mm; TR: 1.46 seconds).
- MEMPRAGE (sessions 1-7, 9-14) Across all sessions (except the full-movie session), we obtained a multi-echo MPRAGE sequence for the purposes of anatomical registration [36]. Within a given scan, four echoes are collected per excitation and combined using root mean square average. This enables the images to be acquired with a higher bandwidth to reduce distortion, while recovering SNR through averaging. The added T2* weighting from the later echoes also helps differentiate dura from brain - matter.
- Diffusional kurtosis imaging (DKI) Leveraging the capabilities of the CMRR multiband imaging sequence, we were able to acquire 64 directions at 2 b-values (1000 and 2000 s/mm2). This enables diffusion kurtosis specific metrics to be calculated from the data in addition to standard DTI metrics and can improve tractography [31].
- Quantitative Relaxometry MRI (Quantitative T1, T2, and Myelin Water Fraction [MWF]): DESPOT1 and DESPOT2 sequences were used to characterize microstructural properties of brain tissue. These innovative acquisition strategies enable quantitation of T1 and T2 relaxation constants, which can be combined to calculate myelin water fraction [37].
- Magnetization Transfer High-resolution T1-weighted structural images were acquired with a FLASH sequence, with and without a saturation RF pulse. The magnetization transfer ratio is calculated from the resulting images, which is purportedly sensitive marker of myelination [34].

## DATA RECORDS

### Data Privacy

The HBN-SSI data are being shared via the 1000 Functional Connectomes Project and its International Neuroimaging Data-sharing Initiative (FCP/INDI) [38]. Prior to sharing, all imaging data were fully de-identified by removing all personally identifying information (as defined by the Health Insurance Portability and Accountability) from the data files, including facial features. All data were visually inspected before release to insure that these procedures worked as expected.

### Distribution for use

#### Imaging Data

All MRI data can be accessed through the Neuroimaging Informatics Tools and Resources Clearinghouse (NITRC; http://fcon_1000.projects.nitrc.org/indi/hbn_ss) and FCP/INDI’s Amazon Web Services public Simple Storage Service (S3) bucket. In both locations, the imaging data is stored in a series of tar files that can be directly downloaded through a HTTP client (e.g., a web browser, Curl or wget). The data is additionally available on S3 as individual NifTI files for each scan, which can be downloaded using a HTTP client or S3 client software such as cyberduck (https://cyberduck.io).

All imaging data are released in the NIfTI file format; they are organized and named according to the brain imaging data structure (BIDS) format [39].

#### Phenotypic Data

Partial phenotypic data will be publically available without any requirements for a data usage agreement. This includes age, sex, handedness, the internal state questionnaire, and the New York Cognition Questionnaire [39]. These data are located in a comma separated value (.csv) file accessible via the HBN-SSI website and are included with the BIDS organized imaging data as tab separate values (TSV) files. The remainder of the phenotypic data (see Table 3), including the PANAS [40] and results from the ADHD Quotient system [41], will be made available to investigators following completion of the HBN Data Usage Agreement (DUA). The HBN DUA is modeled after that of the NKI-Rockland Sample and is intended to prevent against data re-identification; it does not place any constraints on the range of analyses that can be carried out using the shared data, or place requirements for co-authorship. Following submission and execution of the data usage agreement, users can access the phenotypic data through the COINS Data Exchange (an enhanced graphical query tool, which enables users to target and download files in accord with specific search criteria) [42].

**Table 3.** Questionnaires and physical measures collected.

| Questionnaires |  |
| --- | --- |
| Internal State Questionnaire (pre-scan, post-scan) | 3-item self-report questionnaire assessing hunger and thirst. Participants respond on a visual analogue scale ranging from "I am not hungry/thirsty/full at all" to "I have never been more hungry/thirsty/full". Responses are rated from 0-100. Participants complete this questionnaire before and after each scan. |
| New York Cognition Questionnaire (NYC-Q) (post-scan) | 31-item self-report questionnaire that asks participants about the different thoughts and feelings that they may have had while in the MRI scan. Participants are asked to indicate the extent to which their thinking or experience corresponded to each item on a 9-point scale. |
| PANAS (post-scan) | The PANAS-S is a self-administered, 20-item Likert scale assessment that measures degree of positive or negative affect. Users are asked to rate 10 adjectives that measure positive feelings such as joy or pleasure, and 10 adjectives that measure negative feelings, such as anxiety or sadness, on a scale of how closely the adjective describes them in the present moment or over the past week. Items are rated on a five-point scale. |
| Physical Measures |  |
| Vitals | Participant vitals (blood pressure, heart rate, blood glucose level, first day of last menstrual cycle) were collected prior to each scan using standard measurement devices in a laboratory environment. |
| Voice data samples | Audio samples of participant speech were recorded prior to scanning. Each sample consisted of 10 sentences with 5 different implicit emotions (neutral, happy, sad, angry, fearful), 10 non-words, and 2 minutes of free speech. For each sample different sentences were drawn from the same set of emotions; the non-words also differed in each sample but had similar characteristics (ie number of syllables, chunks). Stimuli were presented on a laptop computer screen. Completion of the sample took up to 15 minutes. |
| Quotient ADHD System | Quotient is a computer based task designed to assess three core symptoms of ADHD: hyperactivity, attention and impulsivity. Participants respond to stimuli presented with random timing and random placement on a screen. Completion of the task takes up to 30 minutes. |
| GeneActiv Actimetry Device | Between scanning sessions, participants wore a non-invasive actimetry sensor that recorded heart rate and indices of physical activity and sleep. The device was placed on participants' non-dominant wrist and data was collected at each scanning session. |

## TECHNICAL VALIDATION

### Quality Assessment

Consistent with the established FCP/INDI policy, all completed datasets contributed to HBN-SSI are made available to users regardless of data quality. Justifications for this decision include the lack of consensus within the imaging community on what constitutes good or poor quality data, and the utility of ‘lower quality’ datasets for facilitating the development of artifact correction techniques. For HBN-SSI, the inclusion of datasets with significant artifacts related to factors such as motion are particularly valuable, as it facilitates the evaluation of the impact of such real-world confounds on reliability and reproducibility.

To help users assess data quality, we calculated a variety of quantitative quality metrics from the data using the Preprocessed Connectome Project quality assurance protocol (QAP; http://preprocessed-connectomes-project.org/quality-assessment-protocol). The QAP includes a broad range of quantitative metrics that have been proposed in the imaging literature for assessing data quality [43].

For the structural data, spatial measures include: Signal-to-Noise Ratio (SNR) [44], Contrast-to-Noise Ratio (CNR) [44], Foreground-to-Background Energy Ratio (FBER), Percent artifact voxels (QI1) [45], Spatial smoothness (FWHM) [46], Entropy focus criterion (EFC) [47]. These are shown for different participants in Figure 2. Spatial measures of fMRI data include (Figure 3): EFC, FBER, FWHM, and well as Ghost-to-Signal Ratio (GSR) [48]. Temporal measures of fMRI data include (Figure 4): Mean Frame-wise Displacement (Mean FD) [49], Median Distance Index (Quality) [50], Standardized DVARS (DVARS) [51], Outliers Detection [50], and Global correlation (GCOR) [52]. See Figures 2-4 for a subset of the metrics; the full set of measures are included on the HBN-SSI website in csv format for download. Review of the QAP profiles led us to exclude 3 participants based on excessively high mean FD from the illustrative analyses presented in the next section. Although not a focus of the current work, visual inspection of the figures points to the potential value of this dataset for establishing the reliability of QAP measures. The impact of scan condition on each of the functional QAP measures was examined using a one-way ANOVA. No significant differences were found for any of the measures. In addition, the test-retest reliability of each QAP measure, for each condition, was assessed using the intra-class correlation coefficient (ICC). The results are shown in Table 4.

**Figure 2.**
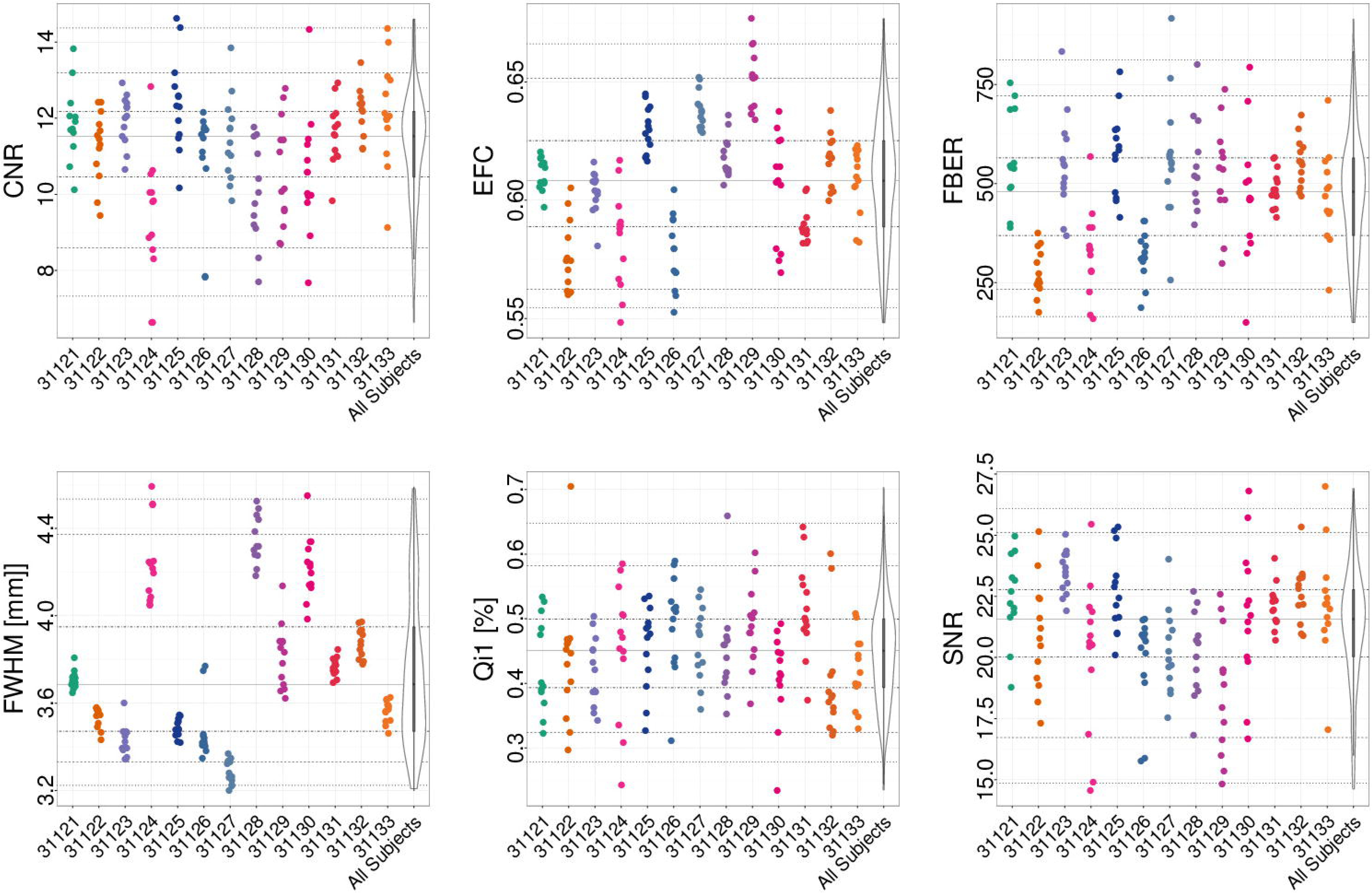
Subset of Quality Assessment Protocol (QAP) spatial anatomical measures for each participant (horizontal axis). Depicted are the following measures: Contrast-to-Noise Ratio (CNR), Signal-to-Noise Ratio (SNR), Entropy focus criterion (EFC). Foreground-to-Background Energy Ratio (FBER), Spatial smoothness (FWHM), Percent artifact voxels (QI1). Each point indicates the measure calculated for an individual scan; for each participant, the data t across scan conditions and sessions are depicted using a single color.

**Figure 3.**
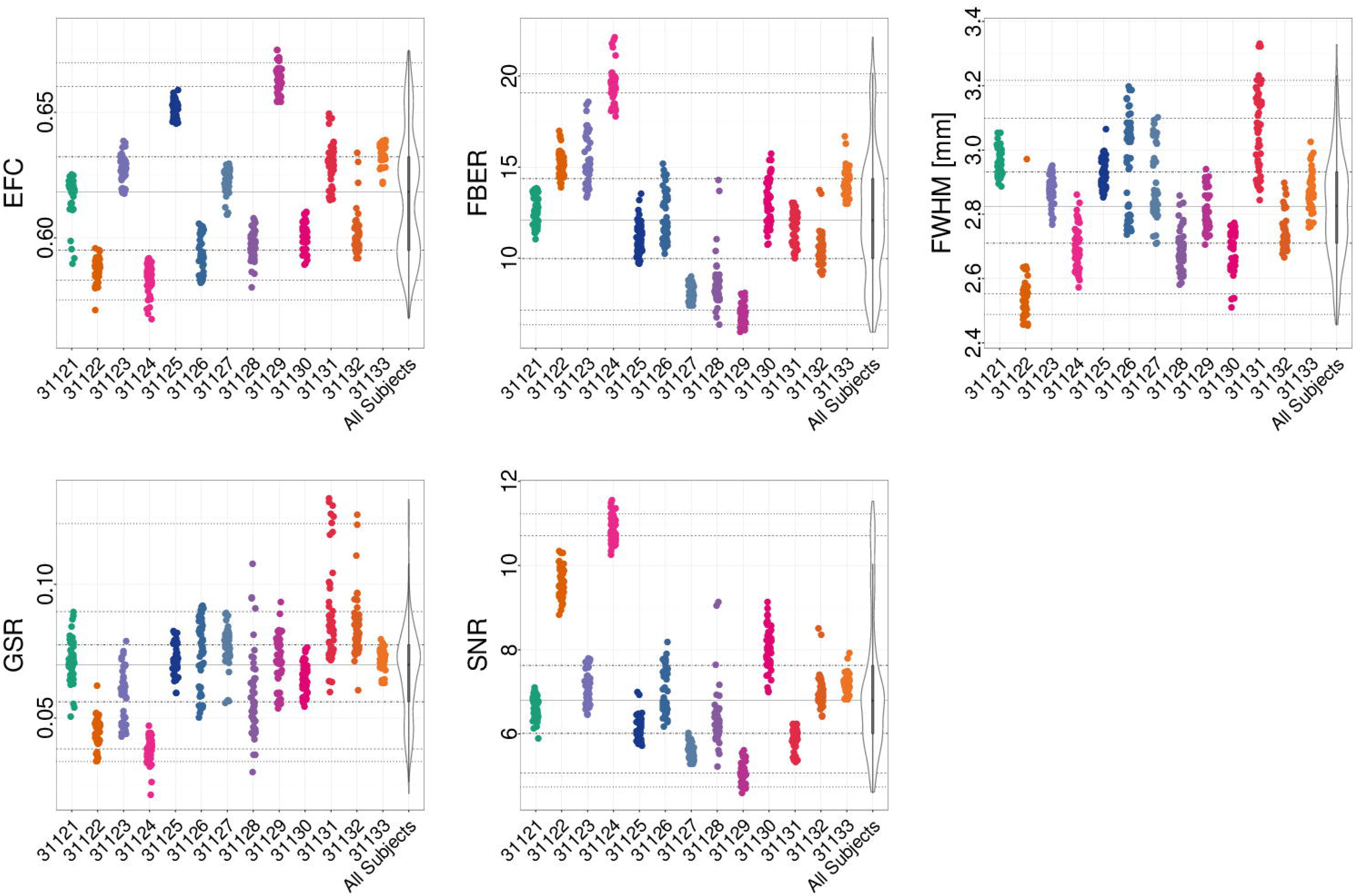
Subset of Quality Assessment Protocol (QAP) spatial functional measures for each participant (horizontal axis). Depicted are the following measures: Ghost to Signal Ratio (GSR), Signal-to-Noise Ratio (SNR), Entropy focus criterion (EFC). Foreground-to-Background Energy Ratio (FBER), spatial smoothness (FWHM). Each point indicates the measure calculated for an individual scan; for each participant, the data t across scan conditions and sessions are depicted using a single color.

**Figure 4.**
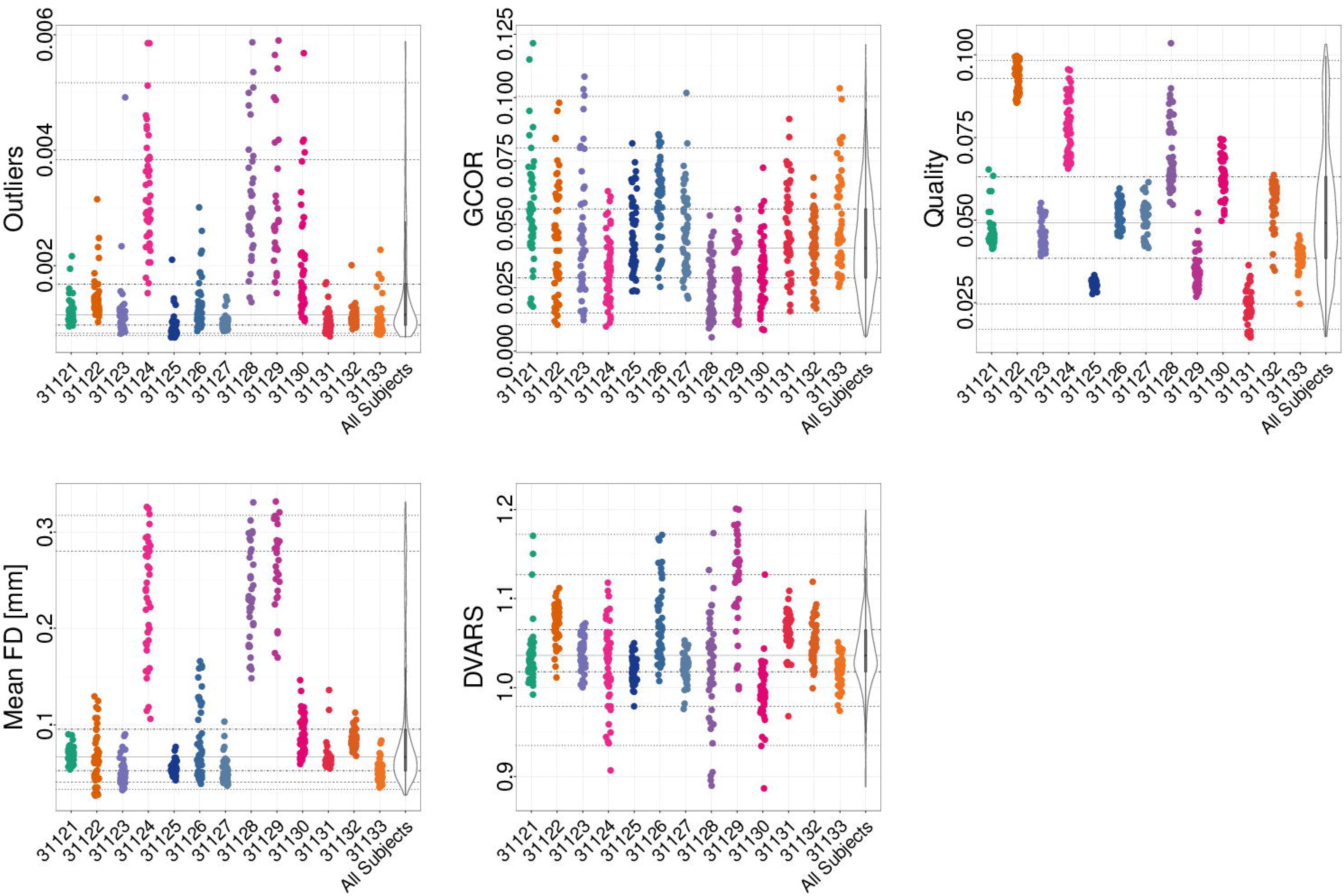
Subset of Quality Assessment Protocol (QAP) temporal functional measures for each participant (horizontal axis). Depicted are the following measures: Outliers Detection (Outliers), Global correlation (GCOR), Quality, Mean Frame-wise Displacement, and Standardized DVARS (DVARS). Each point indicates the measure calculated for an individual scan; for each participant, the data t across scan conditions and sessions are depicted using a single color.

**Table 4.** Test-Retest reliability of quality assurance protocol (QAP) measures, for each scan condition

| Measure | Rest | Inscapes | Movie | Flanker |
| --- | --- | --- | --- | --- |
| EFC | 0.90 | 0.91 | 0.93 | 0.92 |
| FBER | 0.84 | 0.84 | 0.84 | 0.83 |
| FWHM | 0.58 | 0.60 | 0.74 | 0.76 |
| GSR | 0.56 | 0.56 | 0.61 | 0.62 |
| SNR | 0.92 | 0.91 | 0.93 | 0.92 |
| Outliers | 0.08 | 0.18 | 0.06 | 0.50 |
| GCOR | 0.11 | 0.09 | 0.16 | 0.04 |
| Quality | 0.94 | 0.94 | 0.93 | 0.95 |
| Mean FD | 0.30 | 0.39 | 0.40 | 0.68 |
| DVARS | 0.42 | 0.49 | 0.47 | 0.49 |

### FMRI Analyses

A broad range of analyses, including but not limited to evaluations of testretest reliability, can be performed using the present HBN-SSI dataset. Here, we provide a few illustrative analyses to demonstrate the technical validity and utility of these data; they are not intended to be exhaustive.

#### Data preprocessing

Prior to image processing, Freesurfer was used to combine the 12 available MPRAGE images into an MRI robust average image for each individual participant. A non-rigid registration between MPRAGE images and a 2mm MNI brain-only template (FSL’s MNI152_T1_2mm_brain.nii.gz, [53]) was calculated using ANTs [54]. Further anatomical processing included with skull stripping using AFNI’s 3dSkullstrip[55] (to include any voxels in the ventricles incorrectly removed by this utility, the brain mask was augmented using a ventricle mask that was generated by reverse transforming the ventricles included in the MNI atlas into native space for each participant). Next, data was processed using a development version of the open-source, Nipype-based [56]- Configurable Pipeline for the Analysis of Connectomes [1] (C-PAC version 0.4.0, http://fcp-indi.github.io; see https://www.nitrc.org/frs/downloadlink.php/9275 for image preprocessing configuration file).

Following resampling of the functional MRI data to RPI orientation, image preprocessing in C-PAC consisted of the following steps: 1) motion correction, 2) boundary-based registration [57], 3) nuisance variable regression (1st and 2nd order polynomial, 24-regressor model of motion [58], mean WM mask signal, mean CSF mask signal). We then extracted representative time series for each ROI in the CC200 atlas[59] (by averaging within-ROI voxel time series). All possible pairwise correlations were calculated amongst ROI time series to generate a ROI-to-ROI connectivity matrix for each scan in each session for each subject. To facilitate ease of presentation and interpretation for our findings, the connections were sorted by intrinsic connectivity network membership, as defined by Yeo et al. [60].

#### Fingerprinting

Prior work by Finn et al. [22] demonstrated the ability to “fingerprint” individuals based on their functional connectivity matrices. Specifically, they found that the level of correlation between connectivity matrices for data obtained from the same participant on different occasions was markedly higher than that observed for connectivity matrices obtained from different participants; this was true regardless of whether functional connectivity was based on resting state or task activation data. Consistent with their work, we found a dramatically higher degree of spatial correlation between connectivity matrices obtained from the same individual on differing sessions, when compared to differing individuals (Figure 5). Also consistent with their findings, we found this to be true regardless of the scan condition employed.

**Figure 5.**
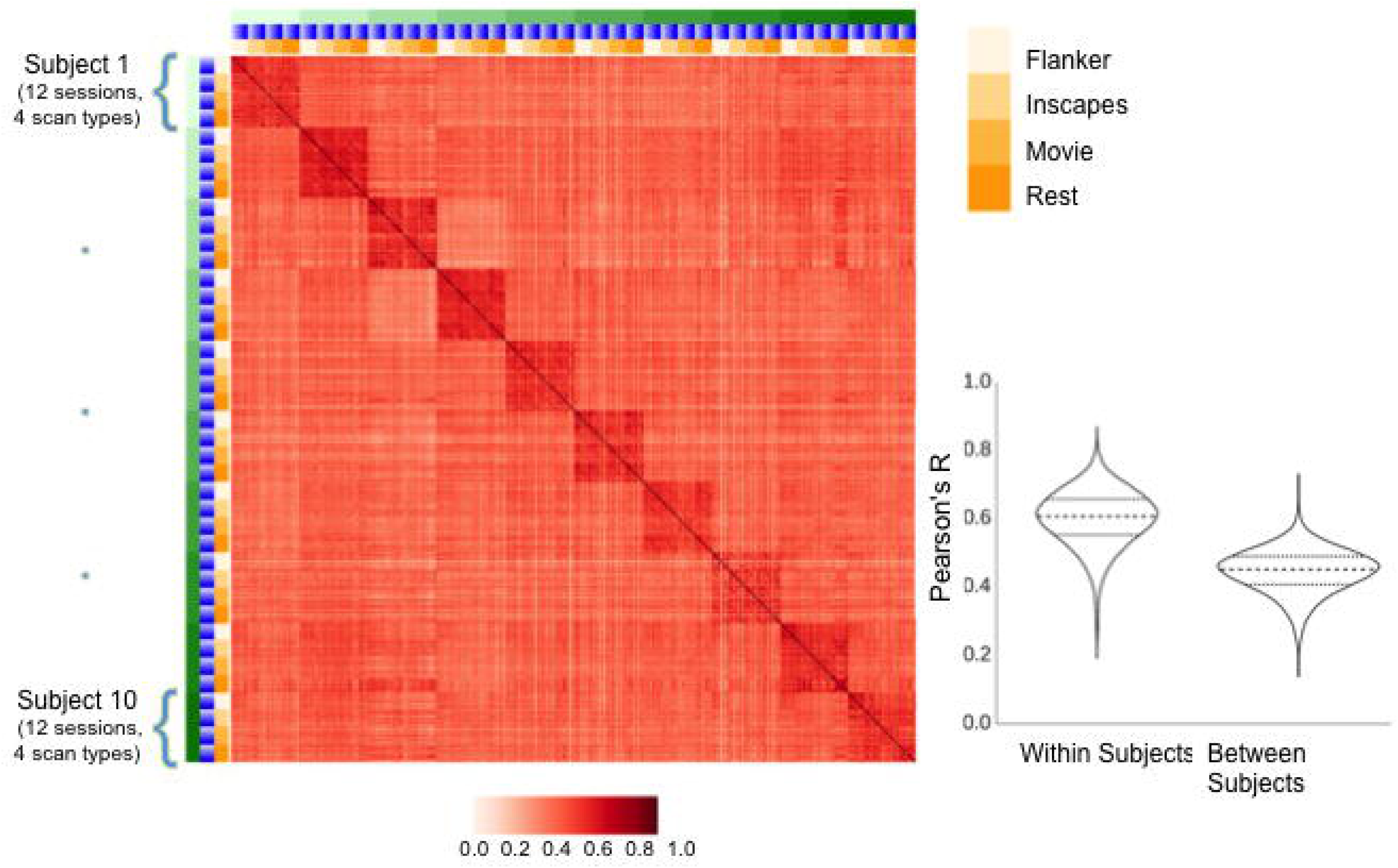
Similarity of full-brain connectivity matrices across participants (green), sessions (blue) and scan conditions (yellow), as measured using Pearson correlation coefficients (red). Also depicted in the bottom right are the distributions of correlation coefficients when comparing scans from the same subject (Within Subject), and scans from different subjects (Between Subject). The distribution of correlation values is also shown (bottom right). On the right column are the values for scans from the same subject, and on the left are scans from different subjects. The median, first, and third quartiles are also depicted with horizontal lines.

#### Connection-Wise Reliability For the Four States

A key question is how much variation among scan conditions (i.e., between-condition reliability) impacts reliability as opposed to between-session reliability (i.e., test-retest reliability). To address this question, we analyzed the 12 sessions obtained for the 10 participants with minimal head motion using a hierarchical Linear Mixed Model (*note:* three subjects were missing the flanker task from one session each; these were treated as missing values in our analyses). The hierarchical LMM allows for the estimation of reliability by providing estimates of variance between participants, across the four conditions (for the same participant) and between sessions within each condition. 
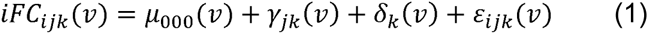

For a given functional connectivity measurement ν, iFC_ijk_(ν) is the modeled intrinsic functional connectivity for the i-th session, for the j-th condition of the k-th participant, taking into account condition and session effects. The equation is composed of an intercept *μ*_000_, a random effect between sessions for the j-th condition of k-th participant *γ*_*jk*_, a random effect for the k-th participant *δ*_*k*_, and an error term *ε*_*ijk*_. *γ*_*jk*_, *δ*_*k*_, and *ε*_*ijk*_, are assumed to be independent, and follow a normal distribution with zero mean. The total variances of iFC can be decomposed into three parts, 1) variance between participants 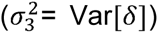, 2) variance between conditions for the same participant 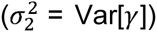, and 3) variance of the residual; indicating variance between sessions 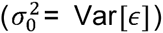. The reliability of the iFC across conditions can be calculated as intra-class correlation coefficients as follows (Figure 6, left):

**Figure 6.**
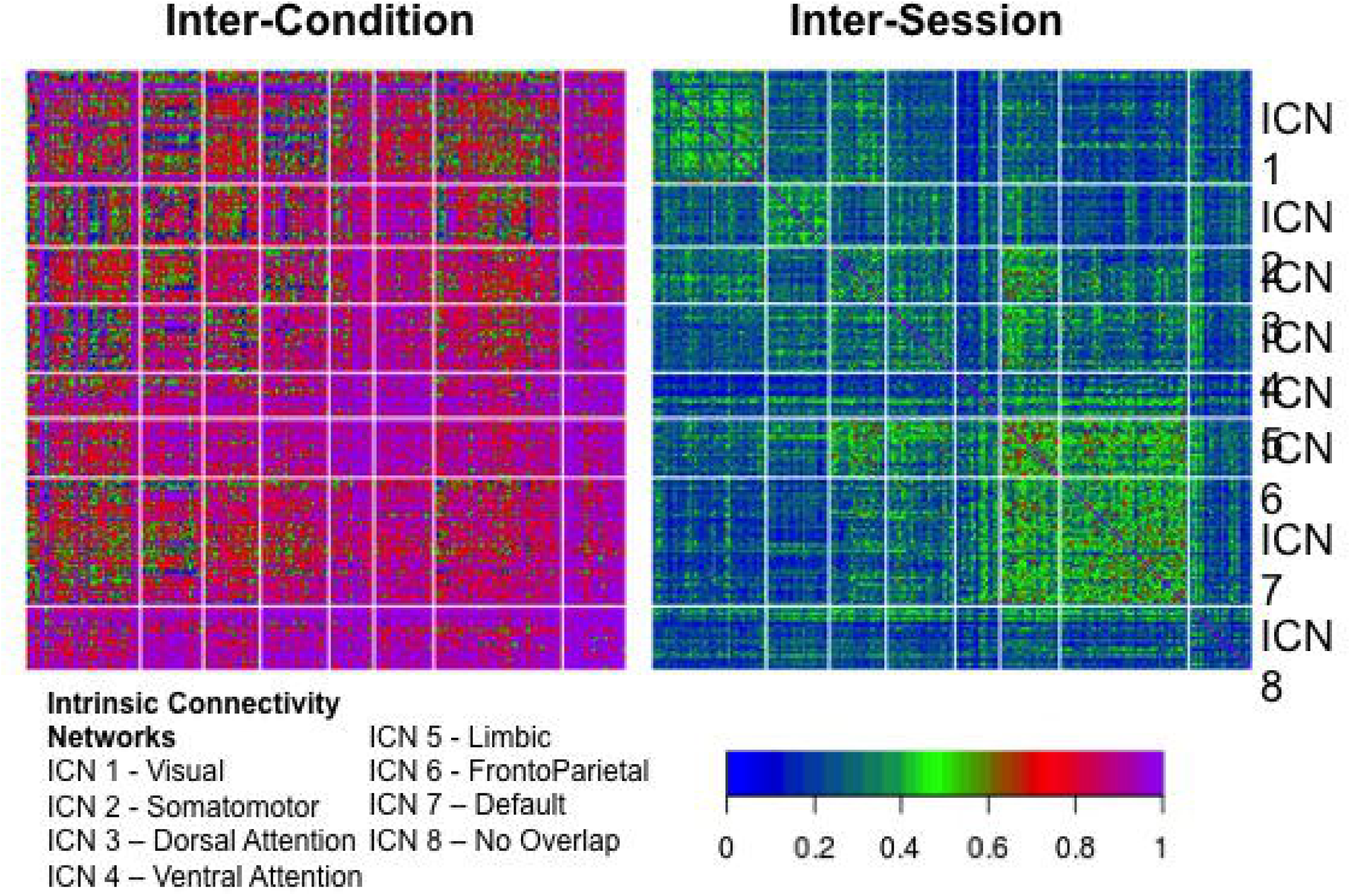
Intraclass correlation coefficients (ICC) quantifying between-condition reliabilities (left) and between-session reliabilities at the connection-level. ICC values were obtained using a hierarchical linear mixed model. These connection-level values are grouped on the vertical and horizontal axes based membership of Intrinsic Connectivity Networks (ICN). No overlap indicates that the voxel did not spatially overlap with any ICN.

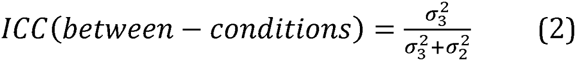
and across sessions as follows Figure 6, right)):

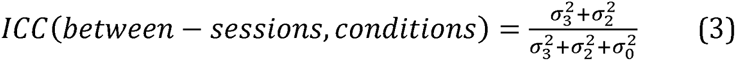

Findings revealed impressively high degree of between-condition reliability for most connections (percentiles: 50^th^: 0.854, 75^th^: 0.955, 95^th^: 1), as opposed to between-session (i.e., test-retest) reliability, which was notably lower (percentiles: 50^th^: 0.270, 75^th^: 0.355, 95^th^: 0.507). Of interest, between-condition reliability tended to be lowest in the visual and somatosensory networks – each of which would be expected to vary in a systematic way across conditions due to differences in visual stimulation (movie > inscapes > flanker > rest) and motor demands (flanker > all other conditions).

Regarding test-retest reliability, follow-up analyses also looked at connection-wise ICC for each of the stimulus/task conditions separately using a linear mixed model (as implemented in R) (see Figure 7), finding similar ranges of ICC scores across conditions, though with some notable differences (e.g., higher ICC for visual network in movies and inscapes; higher frontoparietal ICC’s in flanker task and rest). Additionally, we used image-wise correlation coefficient (I2C2) [61] to look at functional networks and their interactions from a multivariate perspective. As can be seen in Figure 7, a high degree of correspondence was noted between the strength of the reliability for a given network (i.e., I2C2) and the strengths of the reliabilities for the individual edges in the network (i.e., ICC).

**Figure 7.**
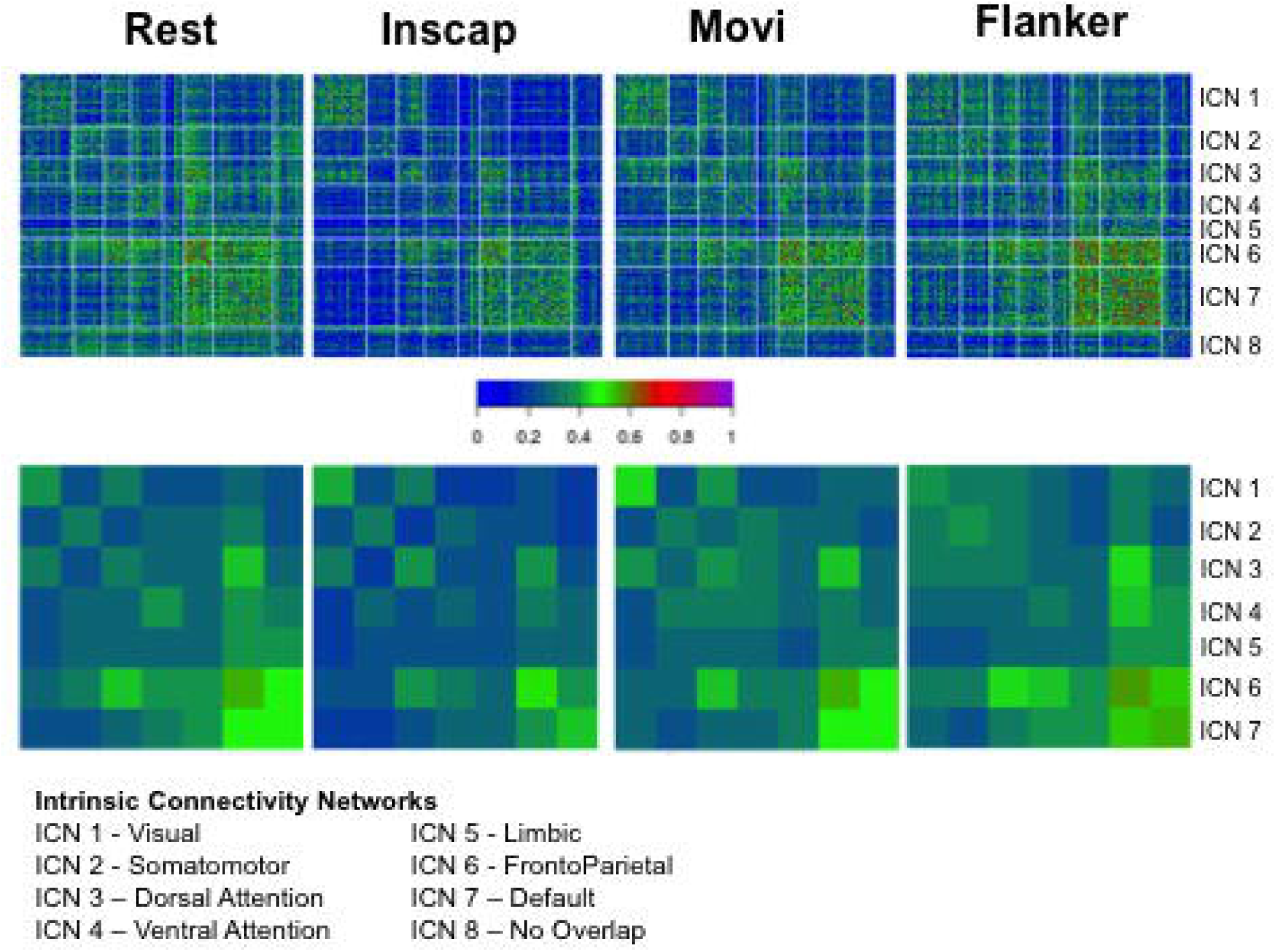
Connection-wise ICC values across all subjects, sessions, and scan conditions (top), as well as network-wise calculations of test-retest reliability carried out using the imagewise intraclass correlation coefficient (I2C2), again across all subjects, sessions and scan conditions (bottom).

Finally, to gain insights into the effects of scan duration on test-retest reliabilities, we repeated ICC and I2C2 analyses using 10, 20 and 30 minutes of scan data across 4 pseudo-sessions (i.e., for 20 minutes, we combined data from 2 sessions; for 30 minutes, we combined data from 3 sessions). Consistent with prior reports, our analyses revealed notable improvement of ICC and I2C2 values with longer scans, particularly when increasing from 20 to 30 minutes (see Figures 8, 9).

**Figure 8.**
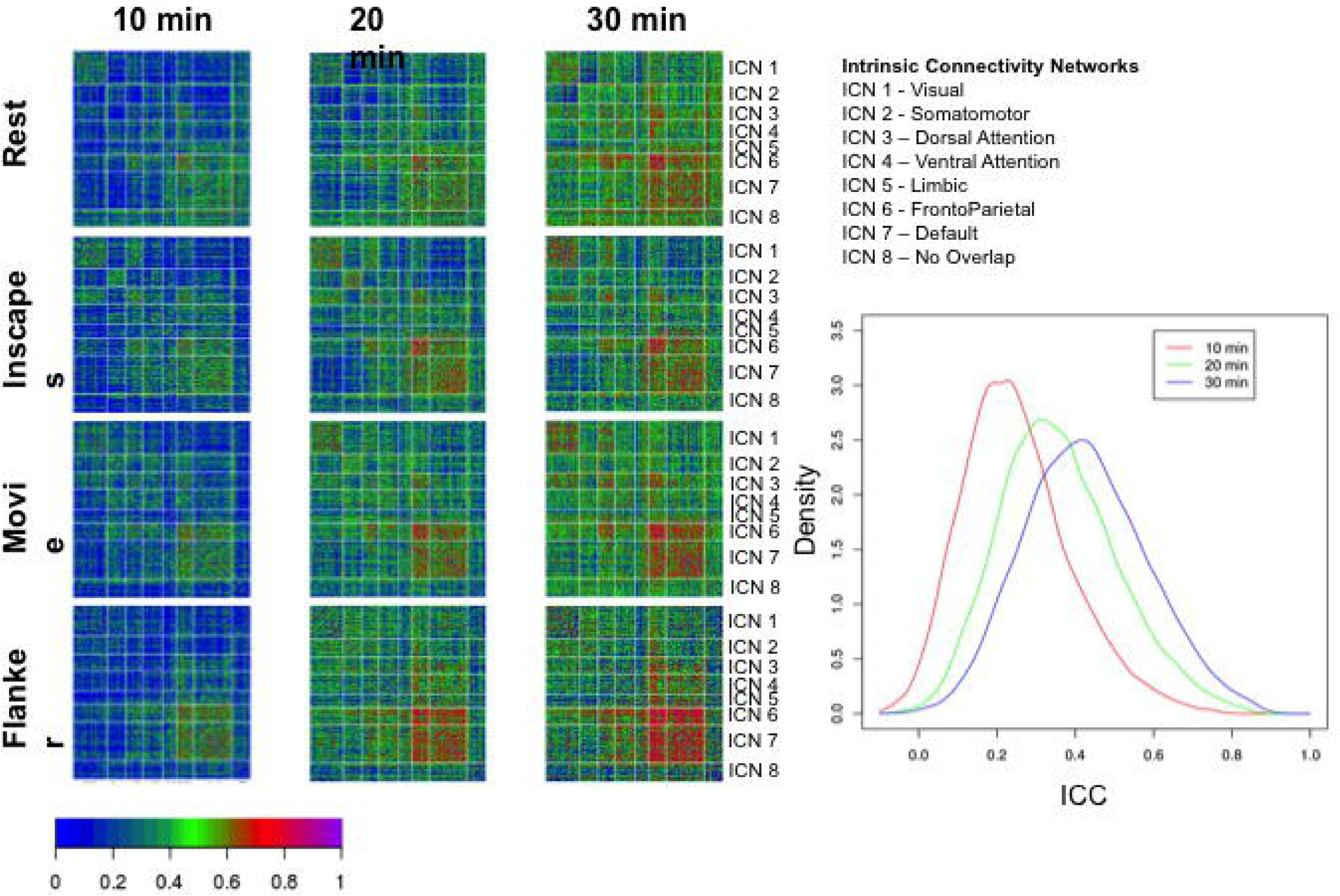
– Impact of scan duration on test-retest reliability at the connection level. We randomly sampled sessions, and concatenated the time series temporally to create pseudosessions of 10, 20 and 30 minutes of data. For each of the pseudosession durations, we depict intraclass correlation coefficients (ICC) obtained for each scan condition. *Note:* across durations, the number of pseduosessions was held constant at four.

**Figure 9.**
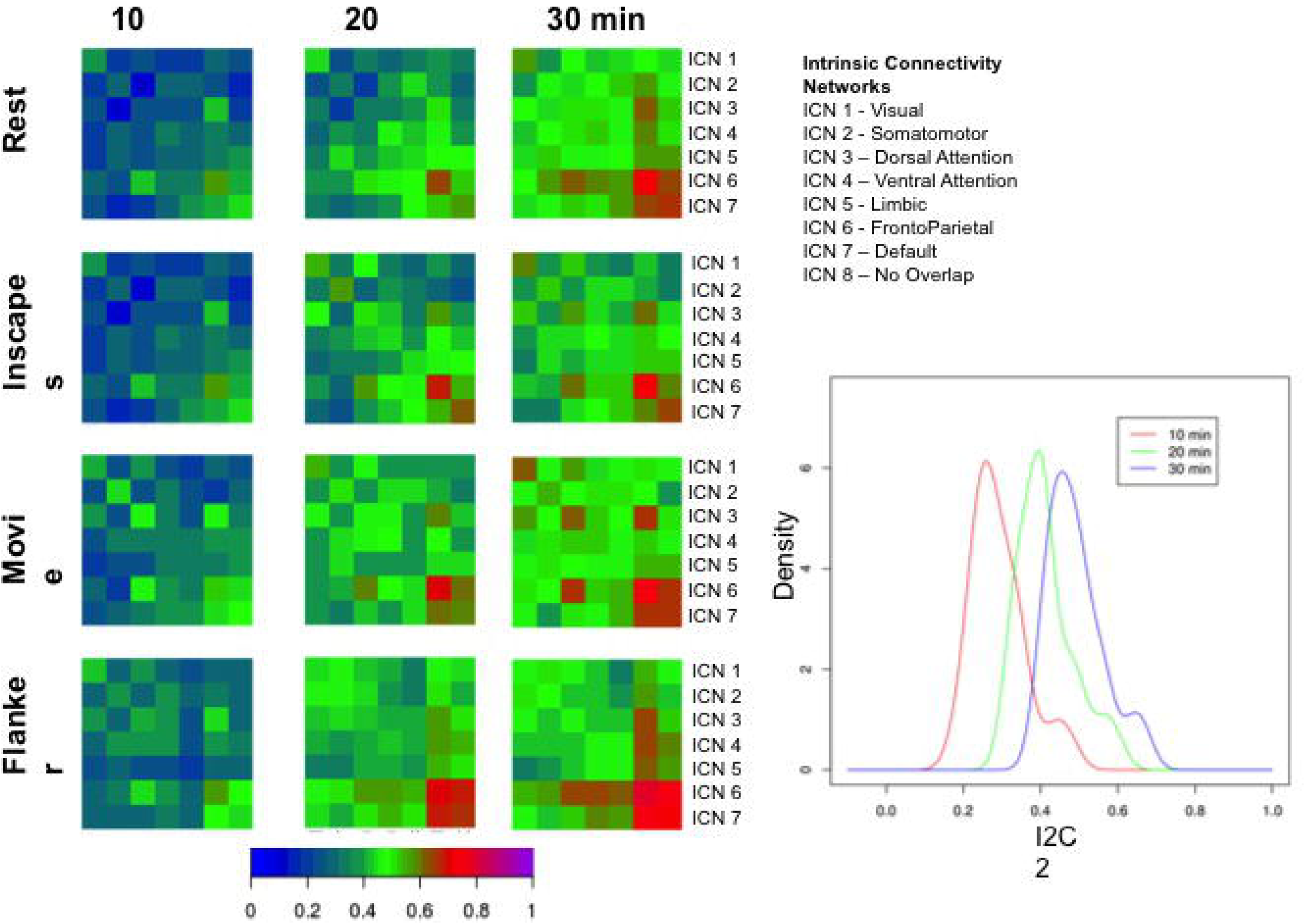
– Impact of scan duration on test-retest reliability at the network level. We randomly sampled sessions to create pseudosessions of 10, 20 and 30 minutes of data. For each of the pseudosession durations, we depict imagewise intraclass correlation coefficients (I2C2) obtained for each scan condition. *Note:* across durations, the number of pseduosessions was held constant at four.

*Concluding Remarks.* These illustrative analyses highlight the value of these data for addressing questions regarding between-condition and between-session reliability. Beyond quantifying reliabilities for connectomic indices, the data available can also be used by investigators to answer questions regarding minimum data requirements (e.g., number of timepoints) and optimal image processing strategies. Finally, it is worth noting that the availability of naturalistic viewing states (Inscapes, movie clips) in the resource will give resting state fMRI-focused investigators an opportunity to explore the added value of these states for calculating intrinsic functional connectivity and more (e.g., exploration of intersubject correlation and inter-subject functional connectivity [23], [62]).

## AVAILABILITY OF SUPPORTING DATA

The HBN-SSI is available at: http://fcon_1000.projects.nitrc.org/indi/hbn_ssi/. The Configurable Pipeline for the Analysis of Connectomes, which was employed to carry out the image processing for the analyses include in the text can be found at https://fcp-indi.github.io; the configuration file containing the settings for C-PAC can be found at https://www.nitrc.org/frs/downloadlink.php/9275.

## LIST OF ABBREVIATIONS

HBN: Healthy Brain Network
SSI: Serial Scanning Initiative
R-fMRI: Resting State Functional Magnetic Resonance Imaging
DKI: Diffusion Kurtosis Imaging
MPRAGE: Magnetization Prepared Rapidly Acquired Gradient Echo
MNI: Montreal Neurological Institute
FSL: FMRIB Software Library
AFNI: Analysis of Functional NeuroImages
ANTs: Advanced Normalization Tools
iFC: Intrinsic Functional Connectivity
ICN: Intrinsic Connectivity Network
CPAC: Configurable Pipeline for Analysis of Connectomes
QAP: Quality Assurance Protocol
SNR: Signal-to-Noise Ratio
CNR: Contrast-to-Noise Ratio
FBER: Foreground-to-Background Energy Ratio
QI1: Percent artifact voxels
FWHM: Full Width Half Maximum
EFC: Entropy focus criterion
GSR: Ghost-to-Signal Ratio
Mean FD: Mean Frame-wise Displacement
GCOR: Global correlation
ICC: Intra-class Correlation Coefficient
I2C2: Image Intra-class Correlation Coefficient

## ETHICS APPROVAL AND CONSENT TO PARTICIPATE

All experimental procedures were performed with approval of the Chesapeake Institutional Review Board and only after informed consent were obtained.

## CONSENT FOR PUBLICATION

All participants consented to have their data shared.

## AUTHOR CONTRIBUTIONS

**Conception and Experimental Design:**

JE, LP, MPM, RCC, SC, SG, TV

**Implementation and Logistics:**

DOC, NVP, RCC, SC, TV

**Data Collection:**

AB, MK, NGV, YO

**Data Informatics:**

DOC, JP, RCC

**Data Analysis:**

DOC, LA, MPM, TX

**Drafting of the Manuscript:**

DOC, MPM, RCC, TX

**Critical Review and Editing of the Manuscript:**

All Authors contributed equally to the critical review and editing of the manuscript.

## AUTHOR DETAILS

1. Child Mind Institute Healthy Brain Network, New York, New York
2. Center for Biomedical Imaging and Neuromodulation, Nathan S. Kline Institute for Psychiatric Research, Orangeburg, New York
3. Yale University, New Haven, Connecticut
4. City College of New York, New York, New York
5. The Graduate Center of the City University of New York, New York, New York
6. Massachusetts Institute of Technology, Cambridge, Massachusetts

